# Deletion of the CTG Expansion in Myotonic Dystrophy Type 1 Reverses *DMPK* Aberrant Methylation in Human Embryonic Stem Cells but not Affected Myoblasts

**DOI:** 10.1101/631457

**Authors:** Shira Yanovsky-Dagan, Ester Bnaya, Manar Abu Diab, Tayma Handal, Fouad Zahdeh, Walther J.A.A. van den Broek, Silvina Epsztejn-Litman, Derick G. Wansink, Rachel Eiges

## Abstract

Myotonic dystrophy type 1 (DM1) results from a CTG repeat expansion in the 3’-UTR of *DMPK*. When the repeat extensively expands, this results in *DMPK* aberrant methylation, reduction in *SIX5* transcription and the development of the congenital form of the disease. To explore whether hypermethylation could be reversed in DM1 embryonic stem cells (hESCs) and patient myoblasts, we monitored methylation levels following removal of the expanded repeat by CRISPR/Cas9-mediated editing. Excision of the repeat in undifferentiated hESCs (CTG2000) resets the locus by abolishing abnormal methylation and H3K9me3 enrichment, and rescues *SIX5* transcription. In contrast, in affected myoblasts methylation levels remain unchanged following deletion of a large expansion (CTG2600). Altogether, this provides evidence for a transition from a reversible to an irreversible heterochromatin state by the DM1 mutation upon cell differentiation. These findings should be taken into account when considering gene correction in congenital DM1 and potentially other epigenetically regulated disorders.

## INTRODUCTION

Myotonic dystrophy type 1 [DM1, (OMIM 160900)] is an autosomal dominant muscular dystrophy that results from a trinucleotide CTG repeat expansion (50 – >3,000 triplets) in the 3’-UTR of the dystrophia myotonica protein kinase gene (*DMPK*) (1, 2). The disease is characterized by myotonia, muscle wasting, cataract, facial and jaw weakness, and frequently by heart conduction defects (3, 4). While DM1 is primarily mediated by a toxic RNA gain-of-function (5), it also features *DMPK* protein deficiency (6, 7), formation of dipeptide repeat inclusions by repeat-associated non-ATG (RAN) translation (8) and down-regulation in transcriptional activity of a downstream gene neighbor, *SIX5* (9–11).

Previous reports, including our own, imply that the transcription levels of *SIX5* are finely tuned by the activity of a narrow, differentially methylated region (DMR) located proximal to the repeat (11–13). Early during development, when the CTG number exceeds 300, this region becomes *de novo* methylated in a way that is dependent on mutation size and is correlated with the cis reduction in *SIX5* transcription. Indeed, *SIX5* reduction has been implicated in several clinical aspects of DM1 pathology including cataracts, male subfertility and heart conduction defects (9, 10, 14–20). In addition, hypermethylation immediately upstream to the expanded CTG repeat was recently shown to be the strongest indicator for the almost exclusive maternal transmission of the congenital and most severe form of the disease (CDM) (21).

How precisely lengthy CTG expansion leads to *de novo* methylation of the 3’-end of *DMPK* remains unknown. One potential mechanism is the binding loss of CTCF immediately upstream to the CTGs, which was suggested to normally counteract *de novo* methylation in that region (22). However data from transgenic mice and hESCs argues against this proposition (12, 19).

In addition, it is still unclear whether hypermethylation, once established, is irreversible in affected cells. Clearly, this issue needs to be addressed when considering the mutation as a potential therapeutic target in affected tissues of CDM. One approach to explore this would be to eliminate the CTG repeat from a heavily methylated allele with a large expansion. This is an alternative therapeutic approach to targeting of expanded, pathogenic *DMPK* RNAs (23–26).

Indeed, the removal of the CTG repeat from *DMPK* using CRISPR/Cas9 has been reported in myoblasts from DM1 patients (27, 28) and trans-differentiated patient fibroblasts (29), DM1 transgenic mice (27), patient-derived induced pluripotent stem cells (iPSCs) and their myogenic cell derivatives (24, 28, 29). These efforts generally restored myogenic capacity and reversed nuclear foci formation, splicing alteration and RNA-binding protein distribution in the corrected cells, providing new therapeutic opportunities for ameliorating disease pathology. Nevertheless, the epigenetic aspects of the disease, which are typically associated with CDM, have not yet been addressed in the repeat-deficient, gene-edited clones.

The aim of this study was to examine whether complete removal of the CTGs from heavily methylated and expanded alleles would reverse the epigenetic modifications that are aberrantly elicited by the mutation in hESCs (12) and patient myoblasts (27). We thereby facilitate dissection of the cause-and-effect relationship between *DMPK* hypermethylation and *SIX5* reduction by uncoupling these two tightly linked molecular events.

## EXPERIMENTAL PROCEDURES

### Cell Culture

The use of DM1-affected embryos, derived from preimplantation genetic diagnosis procedures for hESC line derivation, was performed in compliance with protocols approved by the National Ethics Committee. All cell lines were established at the Shaare Zedek Medical Center (IRB 87/07). Cell line derivation and characterization were carried out as described previously (30). Unaffected hESC line (SZ-13), DM1 hESC line (SZ-DM14, carrying CTG2000 expansion) and all edited cell clones established from these cell lines were cultured in hESC media (knockout DMEM (Gibco) supplemented with 8 ng/mL basic fibroblast growth factor (PeproTech)). HEK-293T cells were grown in Dulbecco’s modified Eagle’s medium, supplemented with 10% fetal bovine serum and 0.5% Penicillin Streptomycin.

DM11 myoblast cell lines and the seven subclones with and without CTG repeat were cultured as described in (27).

### Cloning of gRNAs

Guide RNAs (gRNAs) were designed to target the upstream and downstream regions flanking the CTG repeat using the Zhang lab CRISPR design tool (crispr.mit.edu). Both gRNAs were cloned into the pSpCas9(BB)-2A-GFP (PX458) and pSpCas9(BB)-2A-Puro (PX459) V2.0 from Feng Zhang (Addgene #48138, #62988 respectively) according to the protocol described in (31). In brief, complementary DNA oligomers, containing the gRNA sequence and a BbsI restriction site, were annealed and ligated into PX458 or PX459 with T4 DNA ligase (New England Biolabs). Insertion of the target sequence into the plasmid was verified by DNA sequence using the U6 Forward primer (5’-GAGGGCCTATTTCCCATGATTCC-3’). Target sites upstream and downstream to the repeat (termed 7gRNA: CAGCAGCATTCCCGGCTACA**AGG** and 44gRNA: CAGTTTGCCCATCCACGTCA**GGG**, respectively) with the highest score for cutting efficiency located as close as possible to the repeat, without altering the putative CTCF binding sites, were chosen for further analysis.

### Transfection of gRNAs

HEK-293T cells and hESCs were co-transfected with 1.25 μg of each of the two plasmids containing 7gRNA and 44gRNA, to induce double strand breaks upstream and downstream to the CTG repeat, using TransIT-LT1 transfection reagent (MIRUS) according to the manufacturer’s recommendations. For the HEK-293T, 5×10^5^ cells were seeded per 6 well and co-transfected with two pSpCas9(BB)-2A-GFP (PX458) plasmids containing the gRNAs (7gRNA and 44gRNA) and a GFP expressing plasmid. 48 hours post transfection, cells were harvested for FACS analysis to identify transfection efficiency and DNA was extracted to identify gene editing events by Cas9.

For the hESCs, undifferentiated cells were grown in a feeder-free culture conditions using defined Essential 8 medium combined with vitronectin-(Gibco)-coated surfaces. One day before transfection 1.25×10^5^ hESCs were seeded per 6 well with 10 μM ROCK inhibitor Y-27632 (PeproTech). For generating clonal lines, hESCs were co-transfected with two pSpCas9(BB)-2A-Puro (PX459) V2.0 plasmids containing either 7gRNA or 44gRNA inserts in addition to a puromycin resistance gene. 24 hours post transfection, 0.2 μg/ml of puromycin was added to the media for 48 hours. Feeder cells (mytomycin C-treated mouse embryonic fibroblasts) were laid on the transfected cells to allow expansion of the clonal lines. Single cell puro-resistant clones were isolated manually and re-plated onto feeder cells.

### Screening PCR for Edited Alleles

For validating the targeted deletion of the CTG repeat, genomic DNA was extracted from a small number of cells of each clone and analyzed by Gene Scan analysis. In brief, one colony from each clone was manually isolated and lysed in an alkaline lysis reagent (25 mM NaOH, 0.2 mM EDTA) at 95°C and neutralized by 20 mM Tris buffer (pH 7.4). Lysed cell samples were directly used in PCR and products were analyzed by Gene Scan, to determine the size of the amplified product (5’FAM primer: 5’-CAGCTCCAGTCCTGTGATCC-3’ and 3’ primer: 5’-CACTTTGCGAACCAACGATA-3’). Gene Scan analysis was carried out using ABI 3130 DNA Analyzer.

Successfully targeted clones, as confirmed by Gene Scan analysis, were propagated and further validated for the excision of repeat tract by Sanger sequencing. To distinguish between homozygous null and heterozygous deletions, PCR was carried out across the 7gRNA cutting site (576 bp product, CRISPR cutting site located in the 3’ end of the 3’ primer sequence, primers: 5’-GCTAGGAAGCAGCCAATGAC-3’ and 5’-CATTCCCGGCTACAAGGAC-3’).

### Southern Blot Analysis

Genomic DNA samples (10-21 μg) from candidate clones and unmanipulated controls were digested with SacI and HindIII (Fermentas) restriction endonucleases, separated on 0.8% agarose gels, blotted onto Hybond N+ membranes (Amersham), and hybridized with a PCR Dig-labeled pDM576 probe (primers: 5’-GCTAGGAAGCAGCCAATGAC-3’ and 5’-CATTCCCGGCTACAAGGAC-3’), as previously described (12). To detect aberrant methylation upstream to the repeats, the SacI-HindIII fragments were further processed by parallel restriction with MspI (+MspI) and its methylation-sensitive isoschizomer, HpaII (+HpaII). Detection of DNA fragments was carried out using CDP-Star Chemiluminescent Substrate for Alkaline Phosphatase (Roche).

### Locus-Specific Bisulfite Colony Sequencing

Genomic DNA (2 μg) was modified by bisulfite treatment (EZ DNA methylation kit, Zymo Research) and amplified by FastStart DNA polymerase (Roche). Amplified products using universal bisulfite converted primers that equally detected methylated and unmethylated alleles (primer forward: 5’-TGGTTGTGGGTTAGTGTT-3’ and reverse: 5’-CCCAACAACCTACAACTATTAT-3’, 289 bp PCR product) were TA-cloned, transformed and single colonies were analyzed for CpG methylation by direct sequencing (ABI 3130) using T7 and Sp6 primers.

### Locus-Specific Bisulfite Deep-Sequencing

Genomic DNA (2 μg) was modified by bisulfite treatment (EZ DNA methylation kit, Zymo Research) and amplified by FastStart DNA polymerase (Roche). PCR was carried out using 5’ primer: 5’-TCGTCGGCAGCGTCAGATGTGTATAAGAGACAG<u>TGGTTGTGGGTTAGTGTT</u>-3’, and 3’ primer: 5’-GTCTCGTGGGCTCGGAGATGTGTATAAGAGACAG<u>CCCAACAACCTACAACTATTAT</u>-3’ (*DMPK*-specific sequences are underlined; adaptors for library preparation are at the 5’ end of each primer). Amplified products were quality controlled, for size using the D1000 ScreenTape kit; Agilent Technologies), and for concentration using the Qubit® DNA HS Assay kit (catalog #32854; Invitrogen). Subsequently, the PCR products were bead purified with Agencourt Ampure XP beads (Beckman Coulter) according to the manufacturer’s protocol. Next, amplicons were subjected to a second PrimMax Takara PCR reaction (1st purified PCR DNA (7.5 μl), 2.5 μl N7XX primer (nextera barcode 1), 2.5 ul S5XX primer (nextera barcode 2), 12.5 μl 2x Primstar ReadyMix). PCR program: 98°C for 1 minute followed by 8 cycles of: 98°C for 10 seconds, 55°C for 10 seconds, 72°C for 30 seconds, then 72°C for 5 minutes and finally hold at 10°C. The 2nd PCR was bead purified, quality controlled using the D1000 ScreenTape kit and the Qubit® DNA HS Assay kit, and all resultant libraries were normalized and pooled at 10 nM concentration. Denaturation and dilution of the sequencing pool was performed according to standard Illumina protocol. Ultimately, 1.5 pM of pool (combined with 40% spiked-in) was loaded onto a NextSeq 500 Mid-Output Kit (150 cycles) cartridge (catalog #FC-102-1001; Illumina, San Diego, CA) for high throughput sequencing on a NextSeq 500 instrument (Illumina), with 150 cycle, single-read run conditions.

### RT-PCR

Total RNA was isolated from the cells by TRI reagent extraction, and then 1–2 μg RNA was reverse-transcribed by random hexamer priming and Multi Scribe reverse transcriptase (ABI). Amplification was performed using the primers: 5’-CCAATGTGCACCTCATCAAC-3’ and 5’-GCCGTCTCTGGCTTCAGT-3’ (235 bp product) and Super-Therm Taq DNA polymerase (Jain Biologicals). For allele expression ratio analysis, DNA and cDNA PCR products were purified and sequenced on an ABI 3130XL genetic analyzer. For locus-specific deep-sequencing, RT-PCR was done using the same primers, but with the addition of adaptors for library preparation at the 5’ end of each primer as described for locus-specific bisulfite deep-sequencing.

### T7 Endonuclease I Cleavage Assay to Identify Off-Target Genome Editing Events

Potential off-target sites were identified using the off-target tool of WTSI Genome Editing (WGE) website (32). The T7 endonuclease I assay (T7EI assay) was used according to Van Agtmaal et al. (27). In brief, primers were designed to PCR amplify fragments of approximately 600 bp, which contained the CRISPR target site in the potential off-target sites (Table S2). Approximately 250 ng of the purified products were digested with 10 U T7EI (New England Biolabs) according to the manufacturer’s recommendations. PCR products, before and after the treatment with T7EI, were resolved by electrophoresis on a 1.5% agarose gel.

### Chromatin Immunoprecipitation

Chromatin immunoprecipitation was performed as described (33). In brief, cells were harvested and then fixed, quenched, and washed in 50-ml tubes. Sonication was carried out using Vibra Cell VCX130 with a 3-mm microtip and 30% amplitude in 5 cycles of 10 min. Immunoprecipitation was performed using an anti-H3K9me3 (Abcam Ab8898). Real-time PCR was carried out on an ABI 7900HT instrument (primers are listed in Table S2). ΔΔCt values were normalized according to the negative control to account for differences in precipitation efficiency. *HOXA9* and *APRT* served as a positive and negative controls for H3K9me3, respectively.

### Data Bio-Informatics Analysis

Raw sequence reads were mapped to the human genome (build GRCh37) using Bismark (34). Methylation calls were extracted after duplicate sequences were removed. Data visualization and analysis were performed using custom R and Java scripts. CpG methylation was calculated as the average methylation at each CpG position, and non-CpG methylation was extracted from the Bismark reports. The raw (fastq files) and analysed (text files) data related to deep-sequencing for *DMPK* methylation following bisulfite treatment, and for the measurements of allele expression imbalances (AEI) in *SIX5* in unmanipulated hESCs and gene-edited clones were deposited at the NCBI GEO under accession numbers GSE128904.

## RESULTS

### Deletion of a Large CTG Repeat Expansion from *DMPK* in DM1 hESCs

In an effort to examine whether *DMPK* hypermethylation can be reversed in DM1 hESCs with a completely methylated allele, we aimed to excise the expanded repeat from the mutated gene by a CRISPR/Cas9 approach. Using the gRNA design tool (crispr.mit.edu), we selected a pair of gRNAs to target the sequences that flank the repetitive region that would provide the highest score and would allow to easily distinguish between unmodified and modified alleles. Targeting was directed to the flanking regions, rather than to the repeat itself, to bypass the lack of PAM motifs in the repeat sequence. In addition, this strategy permitted to minimize the high risk for undesired effects at other CTG repeat-containing loci in the genome. To minimize sequence alterations beyond the repeat, we chose for gene editing the nearest pair of gRNAs (7gRNA and 44gRNA, targeting −10 bp and +47 bp relative to the repeat, respectively; Figure 1A). We first validated the efficiency of the selected gRNAs in HEK-293T (5/5 CTGs) cells by PCR, resulting in the amplification of a 189 bp PCR product, if targeting of the gRNA pair was efficient, and/or a 262 bp product, if one/both alleles remained intact (data not shown).

**Figure 1.**
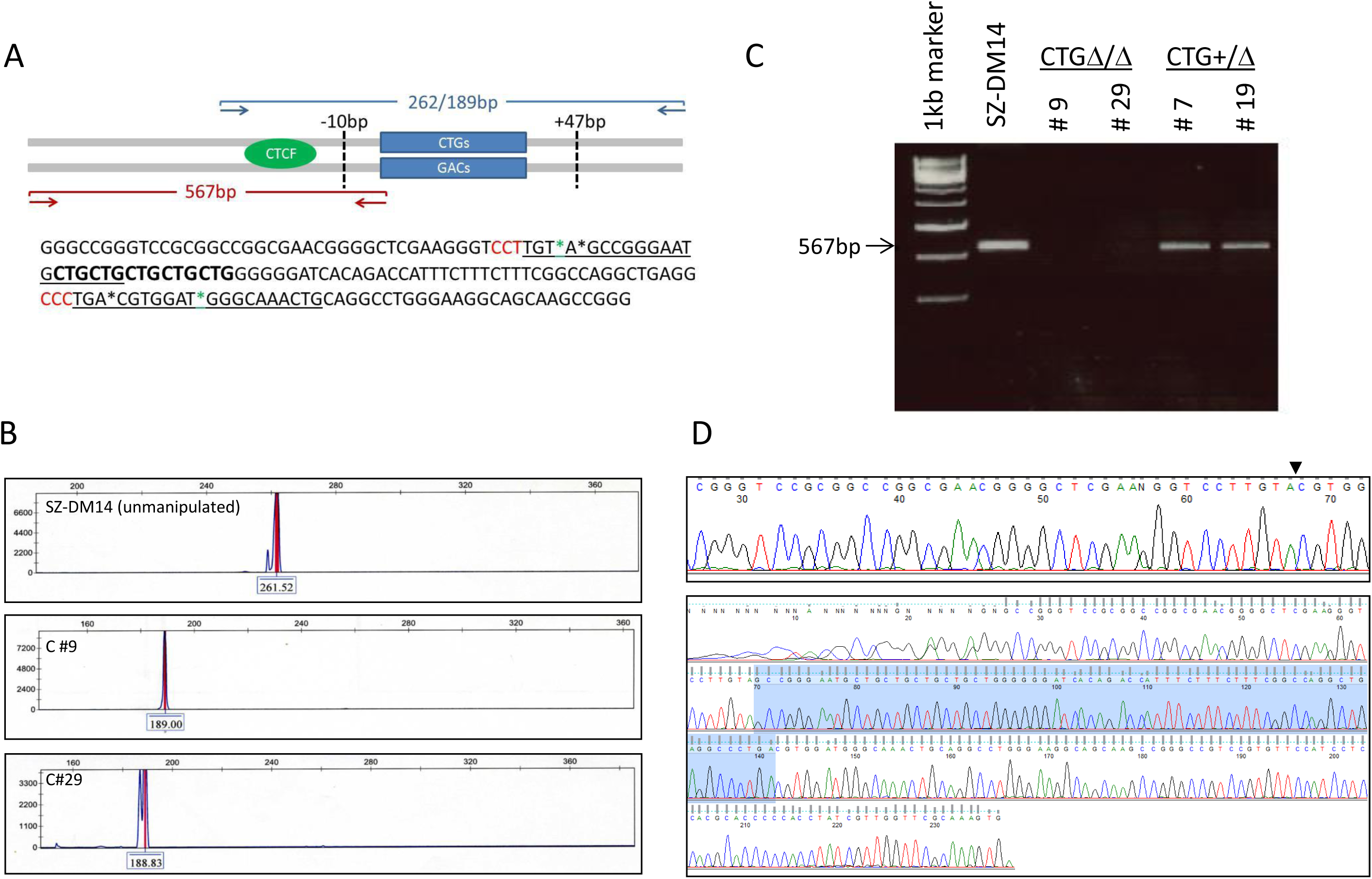
Deletion of Large CTG Repeat in DM1 hESC line. (A) Schematic overview illustrating the CRISPR/Cas9 target sites (dashed vertical lines) relative to the CTG repeat (blue bars), and the PCR primers used for the initial screen using a pair of primers that span the repetitive region (resulting in 262 bp product from intact 5CTG allele or a 189 bp product if targeting was successful); or a primer pair that flanks the 7gRNA target site (567 bp product expected if targeting was not efficient). Marked in black and green asterisks predicted breakpoints in hESCs (7gRNA and 44gRNAs) and myoblasts (27), respectively. (B) Gene Scan analysis of PCR products was used to distinguish between hESC clones that were successfully targeted at least in one of the alleles, resulting in a 189 bp fragment due to the loss of a 72 bp product from the normal (represented by a 262 bp product) or expanded (undetected by PCR) alleles. (C) Absence of PCR products (567 bp) indicates homozygousity (clones #9 and #29) but not heterozygousity (clones #7 and #19) for the deletion of the CTG repeat tract in genetically modified hESC clones and unmanipulated cells (SZ-DM14). (D) Validation of CTG repeat excision by DNA sequencing: DNA sequencing of the repeat and flanking regions in SZ-DM14 (CTG2000) prior (unmanipulated, bottom panel) and following (clone #9, top panel) to gene editing. The site at which the double strand breaks are fused is indicated by a black arrowhead. No indels were found.

Once the targeting potential of the gRNAs was confirmed in HEK-293T cells, we aimed to correct the DM1 mutation by removing the expanded repeat from a hESC line with a heavily methylated (∼100%) CTG2000 expansion (SZ-DM14, 5/2000 CTGs (12)). Considering the particularly low transfection efficiency in hESCs, and the difficulty in growing hESCs as single cell clones, we directed a selection for rare events of gene targeting by transfection with the PX459 Cas9-gRNAs with a puromycin resistance gene. Following a 48-hour antibiotic selection, beginning one day following transfection, we obtained a considerable number (>50) of puro-resistant clones.

As a first step towards the identification of effective editing events, we screened for the expected deletion by Gene Scan analysis of PCR-amplified products from genomic DNA. Successful targeting of the repeat region by Cas9 with 7gRNA and 44gRNA was anticipated to result in a 189 bp fragment if the normal or expanded alleles were properly edited (loss of 57 bp flanking sequence in addition to 5 or ∼2,000 CTG triplets, respectively; Figure 1B). To confirm that the expanded allele was also edited, we PCR assayed the 5’ breakpoint of the 7gRNA. According to this assay, absence of products indicated complete deletion of the CTG repeat from normal as well as expanded chromosomes (Figure 1C). Selected clones exhibiting elimination of the repeat were further validated by Southern blot analysis, leaving no doubt regarding the successful editing of both alleles (Figure S1). DNA sequencing further demonstrated accurate repair of the double strand breaks after repeat excision (Figure 1D).

Altogether, we achieved to establish 27 puro-resistant clones (see Table S1) of which 18 were found to be targeted by either or both Gene Scan and Southern assays. Of those, two were completely *DMPK* repeat deficient with no indels (Δ/(Δ) in normal and expanded alleles (7.4%, clones #9 and #29). In the remaining clones, targeting was less efficient and led to a more complex and unpredictable result. While in most edited clones (12/16) PCR revealed the presence of the normal, jointly with repeat-deficient allele (262 bp and 189 bp fragments), Southern blot analysis indicated either no change in expansion size (#12, #19, #22) or a complete deletion of the CTG tract from the normal allele in addition to CTG tract shortening from the mutant allele (#7 and #13). In three clones (#7, #10 and #26) Southern blot indicated partial targeting resulting in the shorter than expected expanded allele (ranging from 700 to 1400 CTGs). This is most likely due to deletions induced by incorrect repair through non-homologous end joining (NHEJ). The apparent inconsistency in the results between the PCR and the Southern blot assays is most likely due to the difference in sensitivity and quantification ability of these methods. While the PCR test is extremely sensitive and may incorrectly overrepresent rare editing events, the Southern blot (which is much less sensitive and provides rough estimates) records only frequent events of gene targeting (Table S1).

To assess the activity of CRISPR/Cas9 plasmids with 7gRNA or with 44gRNA towards potential off-target sites elsewhere in the genome, we used the off-target tool of WTSI Genome Editing (WGE) website. Potential sites were identified based on similarity to the pair of selected gRNA sequences. Indels in the loci of these off-targets were surveyed by the T7EI assay. Within these loci no indels could be observed, indicating that no cleavage activity outside of the *DMPK* locus occurred with 7gRNA or the 44gRNA as determined by this crude assay (Figure S2).

### Expanded Repeat Excision Eliminates *DMPK* Aberrant Methylation Upstream to the Repeat in Mutant hESCs

To examine whether the elimination of the expanded repeat had reversed aberrant methylation upstream to the CTGs (region E in (12)), we measured DNA methylation levels before and after repeat excision by bisulfite DNA colony sequencing. This is to explore whether aberrant methylation is reversible once the mutation is removed or is preserved by epigenetic memory for many cell generations. Methylation levels were measured 9 passages after gene manipulation by colony bisulfite sequencing (26 CpG sites) 650 bp upstream away from the CTG repeat. This region was formerly identified to be part of the DMR, and becomes hypermethylated when the repeats expands beyond 300 triplets in a way that depends on expansion size in hESCs (12). Using this experimental approach provided clear evidence for a widespread event of demethylation from 55% (unmanipulated parental cells, corresponding with 100% methylation on the mutant allele) to 0% in both of the Δ/Δ CTG-deficient clones (#9 and #29), as well as unaffected hESC line control (SZ-13) (Figure 2).

**Figure 2.**
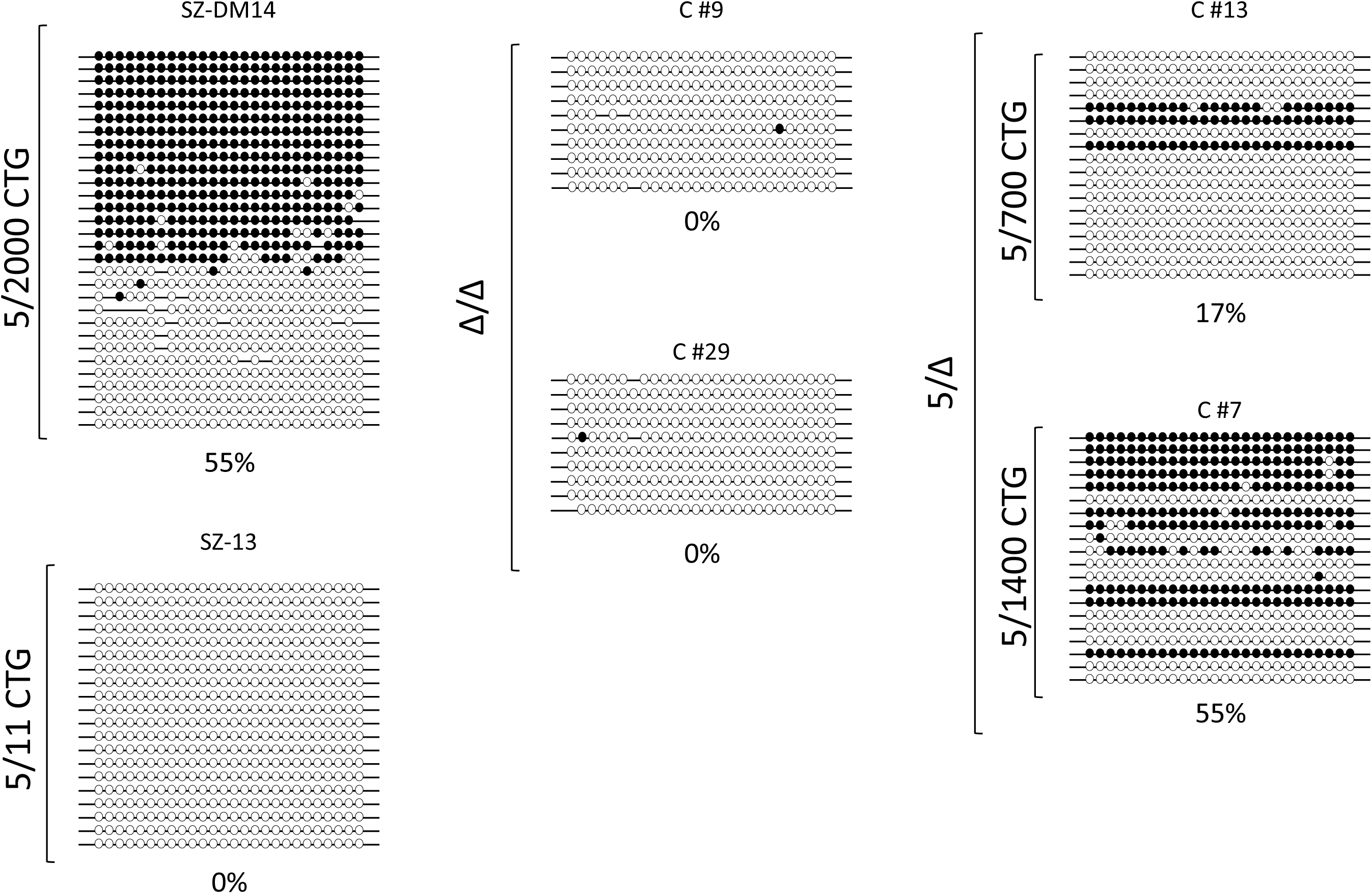
Loss of *DMPK* Hypermethylation by CTG Repeat Excision in Mutant hESCs. Colony DNA bisulfite sequencing of the DMR (region E, 488-777 bp upstream of the repeat, 26 CpG sites) in unaffected with 5CTG/11CTG alleles (SZ-13, methylation levels of 0%), unmanipulated hESC with a heavily methylated CTG2000 expansion (SZ-DM14, methylation levels of 55%,), a pair of gene-edited CTG-deficient ((Δ/Δ) clones (#9 and #29, methylation levels of 1-2%,) and two DM1 hESC clones with 1300 CTGs (55%, #7) and 700 and less CTGs (17%, #13). Full circles correspond to methylated CpGs while empty circles represent unmethylated CpGs.

This observation was further validated by locus-specific deep-sequencing in an overlapping region (15 CpGs, −759 bp to −631 bp relative to the repeat) in the parental cell line and Δ/Δ clones (accession number GSE128904). In addition, different patterns were observed in other clones, where methylation levels were either unchanged (clone #7) or reduced (clone #13) by the failure to remove any or just part of the repeat (1300 copies in #7, 700 copies and less in #13) in a significant number of cells (Figure 2 and Table S1). Altogether, we infer from the methylation analysis at the DMR in the repeat-targeted clones, that *DMPK* hypermethylation in DM1 is dependent on the continuous presence of the expansion mutation in undifferentiated hESCs.

### CTG Repeat Excision Abolishes Heterochromatin in DM1 hESCs

To explore whether hypomethylation is coupled to the loss of heterochromatin, we analyzed the enrichment for H3K9me3 (repressive histone modification) immediately upstream to the repeat by chromatin immunoprecipitation (ChIP). While significant enrichments were obtained in unmanipulated parental hESCs, no enrichments for H3K9me3 could be observed in unaffected hESCs (SZ-13, CTG5/CTG11) or Δ/Δ repeat-deficient hESC clones (#9 and #29) (Figure 3). Taken together, this suggests that repeat removal alters the epigenetic status of the locus in a way that prevents heterochromatin to be re-established nearby the repeat.

**Figure 3.**
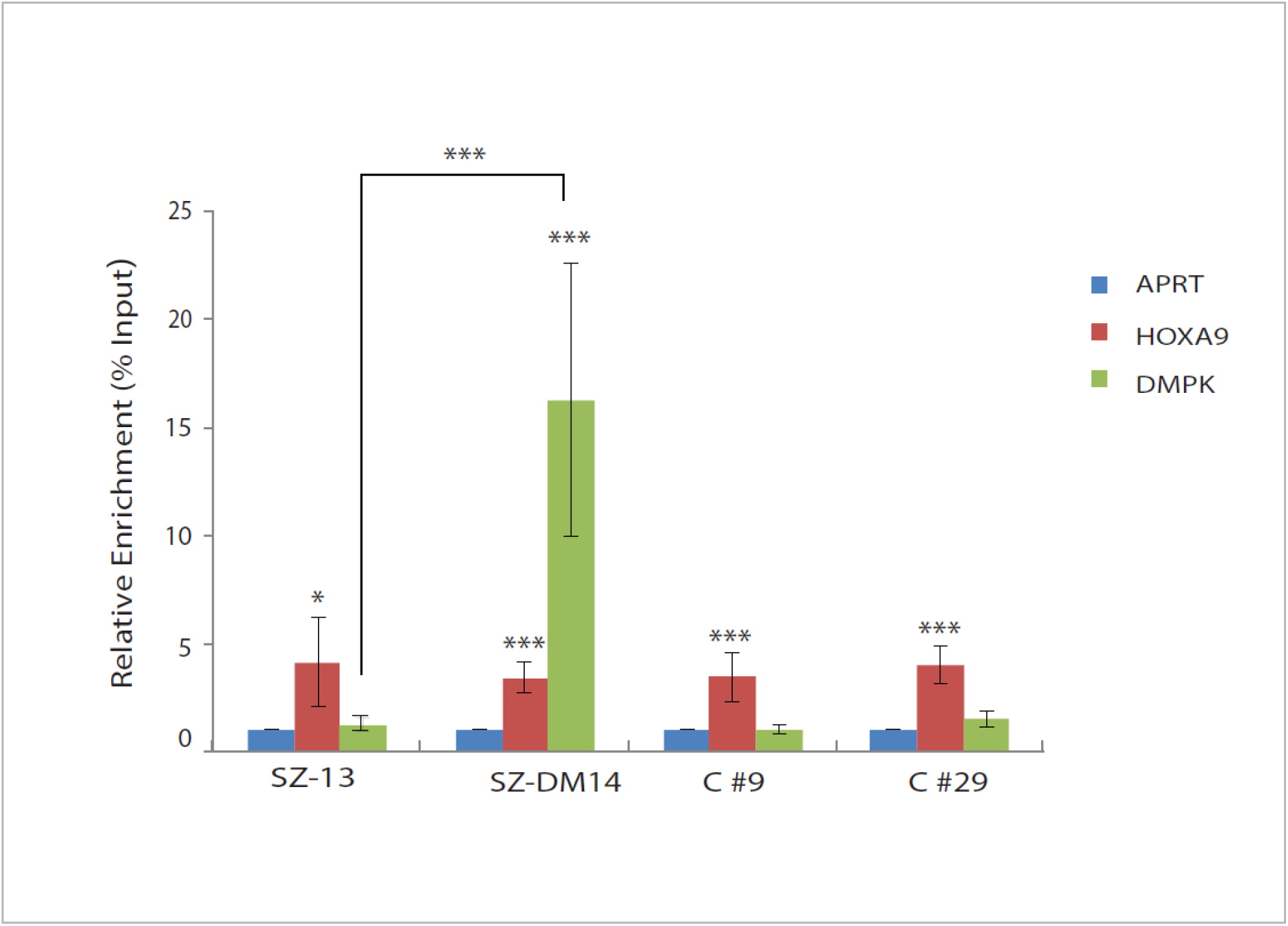
Loss of H3K9me3 at the DM1 Locus after expanded CTG Repeat Excision. Real-time PCR ChIP analysis for H3K9me3 in unaffected (SZ-13), parental DM1 affected hESC line (SZ-DM14, CTG2000), and isogenic CTG-deficient homozygote clones (C#9 and C#29). *APRT* and *HOXA9* were used as negative and positive controls, respectively. Negative controls were set to one. The data in each panel represents an average of 3-4 independent ChIP experiments. Error bars represent standard errors (paired t test, *p < 0.05, ***p < 0.001).

### CTG Repeat Excision from DM1 hESCs Restores *SIX5* Haploinsufficiency

To address the mechanistic association between aberrant methylation levels and the decrease in *SIX5* expression, we took advantage of an informative SNP within exon 3 of *SIX5* (rs2014377) to associate allele-specific gene expression with methylation in the parental hESC line, before and after gene editing, as well as in a genomic DNA control (expected ratio of 1:1 in genomic DNA from SZ-DM14 or any other cell type). We utilized this informative site to identify expression imbalances (AEIs) by precisely measuring the expression ratio between the alleles via locus-specific RT-PCR deep-sequencing (accession number GSE128904). This approach clearly illustrated allele imbalance in *SIX5* transcription in unmanipulated cells, a difference of 0.43 in favor of the normal allele. This contrasts with the CTG-deficient clones, where allele imbalances were reversed, resulting in 1.2 to 1.3 differences in favor of the alleles from which the expansion was removed (Figure 4). We conclude that demethylation upstream to where the expanded repeat was located before it was excised leads to the correction of *SIX5* haploinsufficiency in DM1 cells, as would be expected if the methylation in the DMR would play a role in fine tuning the expression of *SIX5* transcription *in cis*.

**Figure 4.**
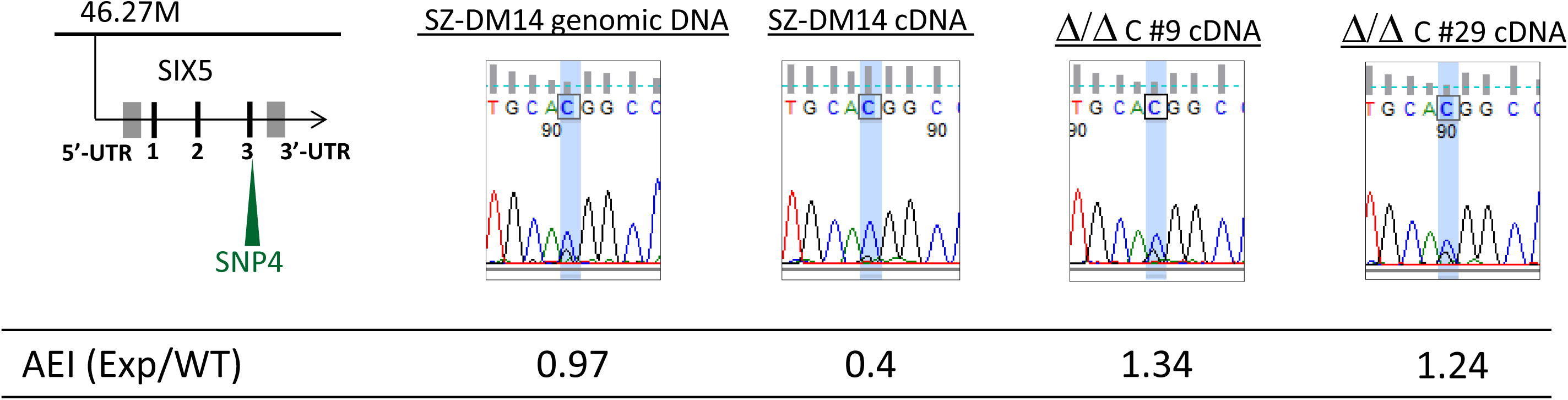
Correction of *SIX5* Haploinsufficiency. Identification of *SIX5* allelic expression imbalances (AEI) by locus-specific DNA deep-sequencing (expanded/normal type) using an informative SNP (rs2014377, designated as SNP4 in (12)) positioned in exon 3 of *SIX5* (12). Results are presented for cDNA samples from unmanipulated DM1 hESCs (SZ-DM14) as well as gene-edited Δ/Δ clones (#9 and #29). Genomic DNA was used as a control for the sensitivity of the assay, presenting nearly 1:1 ratio between the alleles, as expected.

### CTG Repeat Excision Does Not Abolish *DMPK* Aberrant Methylation in Patient Myoblasts

Finally, to explore the relevance of our findings regarding *DMPK* demethylation by repeat removal in affected tissues, we took advantage of previously established patient-derived repeat-deficient myoblasts (27). Considering that the CTG2600 repeat in those cells was targeted with nearly the same pair of gRNAs as for the hESCs (see Figure 1A, marked by green asterisks), these cells provided a good opportunity to compare the effect of expanded repeat excision on the methylation status of the DM1 locus between myoblasts and undifferentiated hESCs. Analysis of DNA methylation levels in myoblast clones with and without CTG2600 repeat was carried out precisely as described for the hESCs (colony bisulfite sequencing 650 bp away from the repeat, 26 CpG sites) after at least 20 population cell doublings. In this case, however, we utilized a non-CpG informative SNP within the DMR to perform allele-specific methylation analysis (rs635299, see also (12)). This allowed us to easily distinguish between normal (CTG13; variant G) and expanded (CTG2600; variant T) alleles during methylation analysis. This approach demonstrated that methylation levels remained unchanged after the complete deletion of the CTG repeat from *DMPK* in three homozygote (Δ/Δ) and one heterozygote (13/Δ) myoblast clones, presenting levels of 100% on the background of the mutant allele (T variant) (Figure 5 and S3). This was in addition to no change in methylation levels of three additional independent clones, where gene editing was inefficient and failed to remove the CTG repeats from either allele (CTG13/CTG2600). Altogether, we infer from this analysis that the removal of the expansion in affected myoblasts cannot reset the normal epigenetic status of the DM1 locus once heterochromatin is induced by a large repeat expansion.

**Figure 5.**
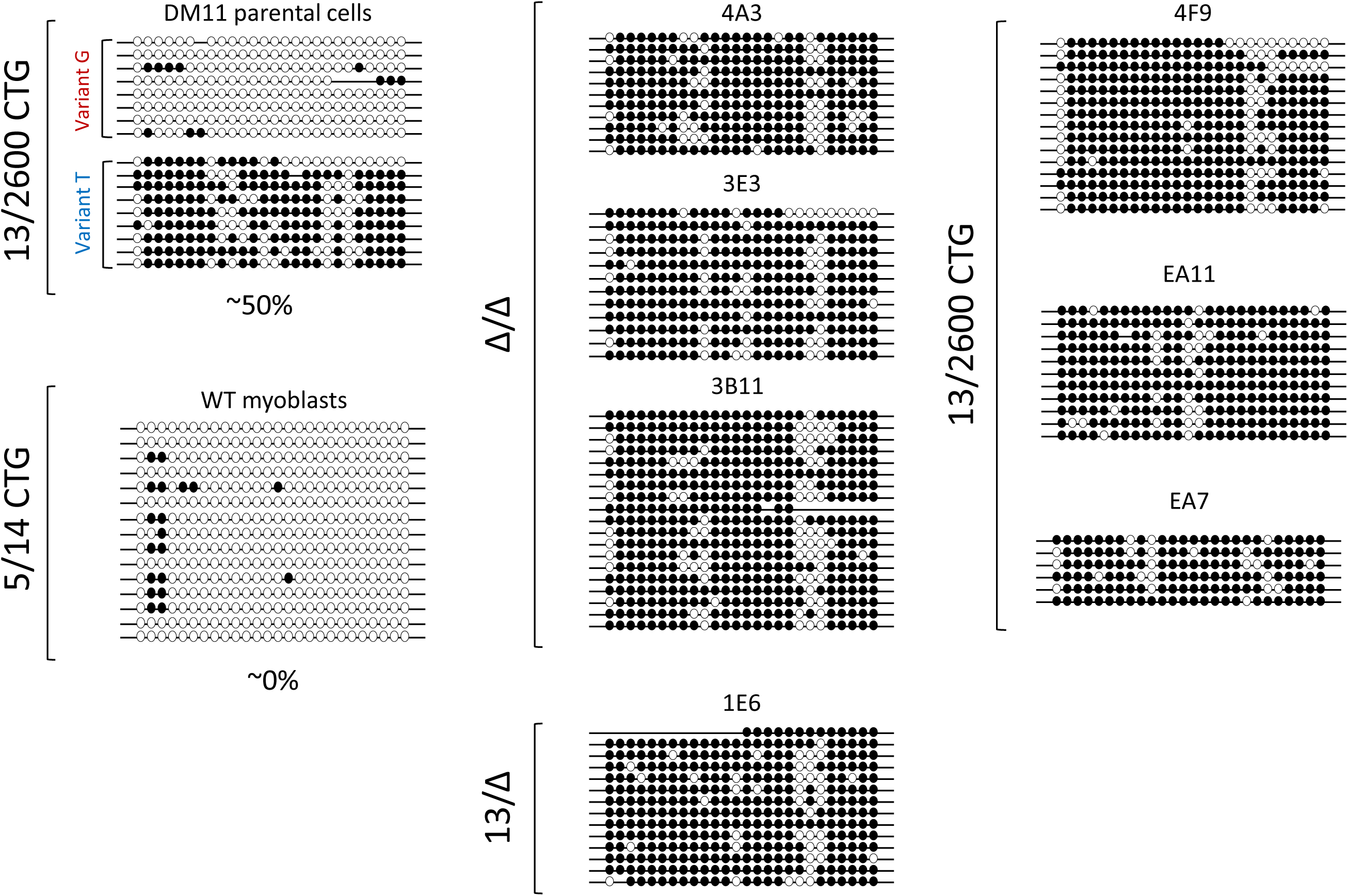
CTG Excision in Affected Myoblasts Has No Effect on *DMPK* Hypermethylation Levels. Allele-specific colony DNA bisulfite sequencing of the DMR (488-777 bp upstream of the repeat, 26 CpG sites) before and after repeat excision (after >20 population cell doublings) from affected myoblasts with normal and expanded alleles (13/2600), three completely CTG-deficient (Δ/Δ), one heterozygote for the deletion on the background of the mutant allele (13/Δ), three unsuccessfully manipulated clones bearing intact alleles (13/2600), and an independent control myoblast cell line (5/14) (27). A non-CpG informative SNP within the DMR (rs635299, see also (12)) allowed to distinguish between molecules obtained from the normal (variant G) vs. expanded (variant T) alleles within each cell line/clone during sequencing. Methylation patterns in all molecules associated with variant T are presented for all manipulated clones (Δ/Δ, 13/Δ and 13/2600). Note that, regardless of the presence of the mutation, all molecules were consistently methylated (100%). In no case normal alleles (variant G) were methylated (see Figure S3). Full circles correspond to methylated CpGs while empty circles represent unmethylated CpGs.

## DISCUSSION

We report complete removal of the CTG repeat from the *DMPK* locus in hESCs with a heavily methylated CTG2000 expansion. By targeting Cas9 to 10 bp upstream and 47 bp downstream of the repeat with a pair of gRNAs, we precisely removed the CTGs from the 3’-UTR of *DMPK* while minimizing the risk of off-target effects on the many CTG repetitive sequences spread throughout the genome. This experimental approach resulted in the establishment of a pair of isogenic hESC clones (2/18 targeted clones) that are *DMPK* bi-allelic repeat-deficient (Δ/Δ) following accurate NHEJ repair.

To address the question of whether hypermethylation can be reversed, we took advantage of the *DMPK*-edited mutant hESCs and compared methylation levels at the DMR, 650 bp upstream to the repeat, prior and following repeat removal. We show that methylation levels extensively decline from 55% (corresponding to 100% on the mutant allele in unmanipulated control cells) to practically 0% in both Δ/Δ hESC clones. Demethylation is coupled to the loss of H3K9me3 enrichment, providing evidence for a general change in the chromatin structure of the locus, i.e. a switch from closed (heterochromatin) to open (euchromatin) configuration. This finding implies that the repressive marks that are elicited by the repeat mutation are reversible and need to be re-established after every DNA replication cycle. In other words, it provides evidence that in undifferentiated cells the CTG expansion is not only essential for triggering heterochromatin, but also for maintaining it. Our data regarding the demethylation of *DMPK* by deletion of the CTG expansion is in line with similar experiments concerning fragile X syndrome, where CGG repeat excision re-activated *FMR1* and led to demethylation of the gene, at least in part, in fragile X iPSCs (CGG repeat in 5’-UTR of *FMR1*) (35, 36).

To link abnormal methylation with disease symptoms, we addressed the question of whether demethylation of *DMPK* is coordinated with the rescue of allele reduction in *SIX5* transcription. *SIX5* has been attributed to various aspects of DM1 pathology including cataracts, male subfertility and heart conduction defects (9–11, 18–20). Taking advantage of an informative SNP in the coding region of *SIX5* (rs2014377, Exon 3, designated SNP4 in (12)), we demonstrate a three-fold increase in expression from the mutant chromosome, as exhibited by a shift in expression ratio between the alleles (expanded/normal) from 0.4 to 1.24-1.34 in the Δ/Δ hESC clones, as determined by locus-specific RT-PCR deep-sequencing. Consistent with our previous report regarding the potential role of the DMR to enhance *SIX5* transcription in reverse correlation with DNA methylation (12), these results support the proposed mechanistic relationship between CTG expansion, DMR hypermethylation and the reduction in *SIX5* expression in DM1.

While this work was being conducted, we reported on the excision of the CTG repeat from the *DMPK* gene (27) in affected myoblasts (CTG2600) by dual CRISPR/Cas9 approach utilizing a pair of gRNAs that overlap with the gRNAs that were used for targeting the repeat in the hESCs. In line with the findings in hESCs, we completly excised the repeat tract from normal and expanded *DMPK* alleles, without inducing changes in the flanking sequences. In that study we provided evidence for that the removal of the expanded CTGs reversed the formation of toxic RNA inclusions that are typically associated with the RNA aspects of the disease. However, the complete elimination of the repeat did not alter expression, splicing nor translational capacity of *DMPK* in muscle cells of patients (27). Also, no change could be detected in *SIX5* transcription in the edited versus unmanipulated parental muscle cells. A proposed explanation for this was that *SIX5* may already be in a heterochromatic repressed state in these cell types, implying that repeat deletion could not reverse the epigenetic status of the region once established, and suggested that muscle cells may not be the proper cell type to assess cis-acting repeat effects on the transcriptional activity of the *SIX5* promoter.

To address the question whether CTG excision could equally remove methylation in these and potentially other disease-targeted tissues, we monitored for a change in *DMPK* methylation following gene editing in the manipulated myoblasts using precisely the same experimental approach as for the DM1 hESCs. Without exception, we find no change in the methylation levels of the mutant allele following repeat deletion in all the manipulated myoblast clones as exhibited by levels of 100% in all tested molecules on the background of the mutant allele (rs635299, variant T). This includes three Δ/Δ and one 13/Δ clone, and three clones where editing was ineffective. Clearly this illustrates a difference in the chromatin structure of the locus between mutant undifferentiated versus differentiated cells, implying a more permissive state in undifferentiated cells. Moreover, consistent with the fact that repeat deletion was unable to reset the epigenetic status of the locus, the Δ/Δ myoblast clones also provide an exceptional opportunity to uncouple repeat expansion from hypermethylation to unambiguously demonstrate that it is the epigenetic modifications, rather than a direct effect of the mutation itself, that are responsible for *SIX5* haploinsufficiency, as formerly suggested (12).

The discrepancy between both cell types in the epigenetic state of the gene resembles the effect of *Xist* (X inactivation specific transcript) in eliciting X inactivation in mouse XX ESCs, where a shift from reversible to irreversible chromosome inactivation occurs during differentiation (37). Using an inducible *Xist* expression system, it was possible to show that *Xist* is responsible for initiating X chromosome-wide epigenetic gene silencing in undifferentiated and early differentiated mouse ESCs (72 hours). This contrasts with fully differentiated XX cells, where *Xist* is no longer required for the maintenance of the inactive state. However, while in early differentiating cells *Xist*-mediated X inactivation is irreversible once established, in undifferentiated ESCs it is reversible and is dependent on the continuous expression of *Xist*. Interestingly, unlike in terminally differentiated XX cells, reversible X inactivation does not involve a shift from early to late replication timing or to a general change in histone H4 hypoacetylations. Taken together, these experiments provide evidence for a transition from a reversible to an irreversible chromatin state at the level of a whole chromosome in ESCs upon differentiation and propose a two-step course for X inactivation: (1) primary inactivation (reversible), and (2) maintenance of the inactive state (irreversible). It is suggested that the primary inactivation step provides a substrate for “locking up” the inactive state later during cell differentiation/embryo development.

By analogy to *Xist*-mediated epigenetic silencing, a long CTG repeat expansion in the DM1 locus in early embryonic cells is anticipated to provide the initiating step for “locking up” the heterochromatin in that region later in development (i.e. somatic cells of patients) (see Figure 6 for model). Our findings are consistent with the preception that DNA methylation during preimplantation development is sequence specific, while in somatic cells it is not. It would be central to explore the changes in specific epigenetic modifications or/and expression of various chromatin modifying enzymes upon cell differentiation to explain the difference in the reversibility of this process. Clearly, the inability to reset the epigenetic modifications that are typically acquired in DM1 by large expansions merely by removing the repeat should be taken into account when considering the mutation as a potential therapeutic target for CDM in disease-relevant tissues. Beyond DM1, our findings may have broader implications to all disease-causing mutations that are involved with aberrant DNA methylation early during development counting noncoding repeat expansion pathologies like fragile X syndrome and genomic imprinting disorders caused by imprinting center defects.

**Figure 6.**
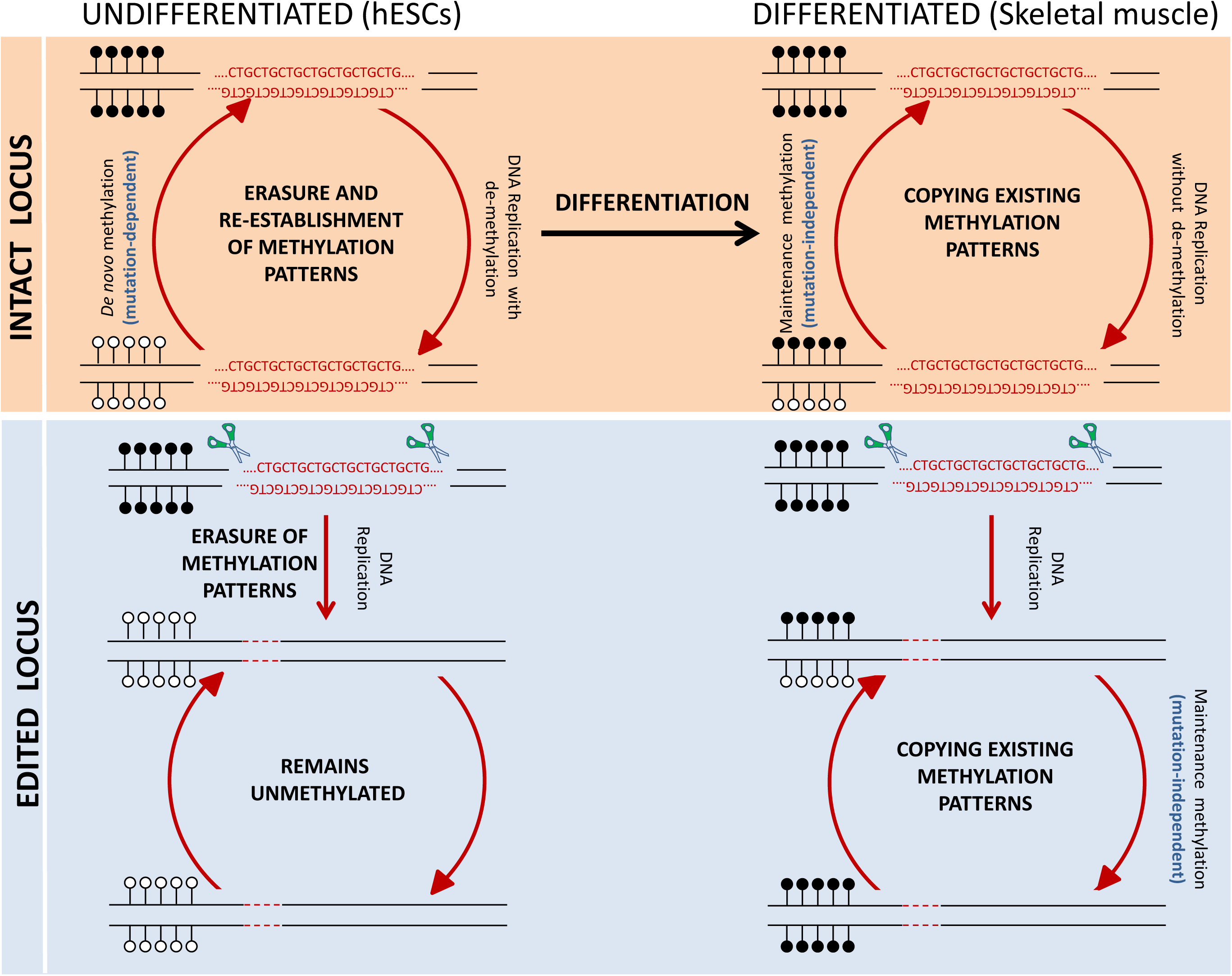
A shift from reversible to irreversible aberrant methylation in DM1 following cell differentiation. *Top panel*: In DM1 undifferentiated embryonic cells (hESCs), aberrant methylation in DM1 is recurrently re-established by *de novo* DNMTs (initiation step) and depends on the presence of the mutation. Once the cells differentiate (Skeletal muscle), aberrant methylation patterns remain unchanged by the activity of maintenance DNMTs (“locking up” step). *Bottom panel*: When the CTG expansion is excised from a heavily methylated allele with a large CTG expansion, it results in the loss of aberrant methylation in undifferentiated cells (reversible). This is contrast to differentiated cells, where excision of the CTGs does not change the epigenetic status of the locus (irreversible). Black lollipops correspond to methylated CpGs while white lollipops represent unmethylated CpGs.

## FIGURE LEGENDS

<u>Figure S1 - Validation of CTG Repeat Deletion in Edited Clones by Southern Blot Analysis</u> Validation of CTG excision by Southern blot analysis in untransfected SZ-DM14 (2000 CTGs) cells and five representative clones after transfection, including #9 and #29 (Δ/Δ). Restriction analysis using this assay allows to distinguish between: normal alleles (∼1.8 kb fragment with 5 CTGs), original expanded alleles (∼8.6 kb fragment with ∼2000 CTGs), completely CTG-deleted allele (∼1.7 kb) and fragments between 1.8 and ∼8.6 kb (representing mutant alleles that have undergone partial deletion by the CRISPR/Cas9 system, designated by asterisk).

<u>Figure S2 - Analysis of CRISPR/Cas9-mediated cleavage directed by 7gRNA and 44gRNA at potential off-target loci.</u>

Cleavage of CRISPR/Cas9 with 7gRNA or 44gRNA at predicted off-target sites (Table S3) was assessed using T7EI assay. DNA fragments containing these putative sites were PCR-amplified on DNA isolated from transfection-positive clones of CRISPR-treated SZ-DM14 (#7 and #13). For each off-target site, we present the edited clones in the absence (left) and presence (right) of T7EI. Potential off-target sites for 7gRNA are presented in the top panel and for 44gRNA in the bottom panel. Note that essentially no cleavage products were observed at the predicted off-target sites.

<u>Figure S3 - CTG Excision in Affected Myoblasts Has No Effect on</u> *<u>DMPK</u>* <u>Hypermethylation Levels</u>

Allele-specific colony DNA bisulfite sequencing of the DMR (488-777 bp upstream of the repeat, 26 CpG sites) before and after repeat excision from affected myoblasts in parental cell line with a normal (CTG13) and expanded (CTG2600) alleles (13/2600), 3 completely CTG-deficient (Δ/Δ), one heterozygote for the deletion on the background of the mutant allele (13/Δ), 3 unsuccessfully manipulated clones (13/2600), and a control cell line (5/14). Taking advantage of a non-CpG informative SNP within the DMR allowed to distinguish between molecules obtained from the normal (variant G) from expanded (variant T) alleles within each cell line/clone during bisulfite sequencing. Methylation patterns in all molecules associated with variant G are presented for all manipulated clones (Δ/Δ, 13/Δ and 13/2600). Note that, in no case normal alleles (variant G) were methylated. Full circles correspond to methylated CpGs while empty circles represent unmethylated CpGs.

## ACKNOWLEDGEMENTS

We thank Dr. Anat Marom for her assistance with pSpCas9(BB)-2A-GFP (PX458) and pSpCas9(BB)-2A-Puro (PX459) plasmid construction, and Dr. David Zeevi and Prof. Bé Wieringa for critical reading of the manuscript.

This work was partly supported by the the ISRAEL SCIENCE FOUNDATION (711/12, R.E.), a donation from the ABRASBA FOUNDATION [Gindi family, to R.E.], LEGACY HERITAGE BIOMEDICAL PROGRAM OF THE ISRAEL SCIENCE FOUNDATION [1260/16, R.E.], ZonMw (TOP grant NL91212009, to Bé Wieringa and DGW) and the Prinses Beatrix Spierfonds (grant number W.OR16-09, to DGW).

## AUTHOR CONTRIBUTIONS

S.Y.D., E.B., M.A.D., T.H., W.V.D.B and S.E.L. conducted the experiments and analysed the data; F.Z. conducted the bioinformatic analysis and D.G.W. and R.E. designed the experiments and wrote the paper.

