## Supplementary figures and images for "Deletion of the CTG Expansion in Myotonic Dystrophy Type 1 Reverses *DMPK* Aberrant Methylation in Human Embryonic Stem Cells but not Affected Myoblasts"

### Supplemental Figures S1-S3

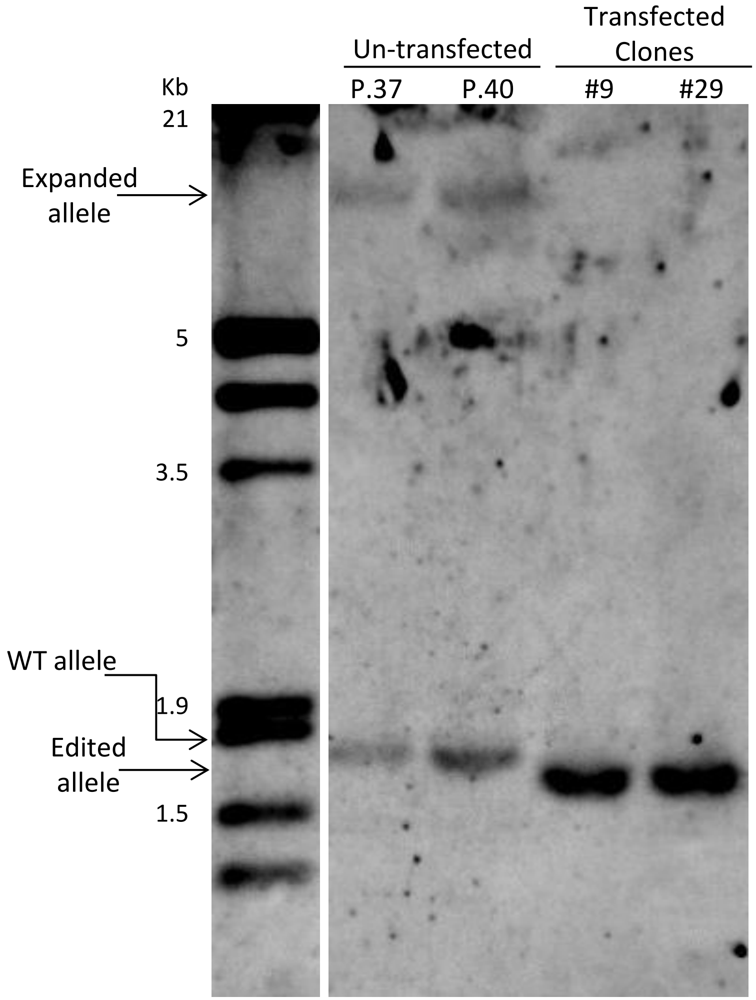

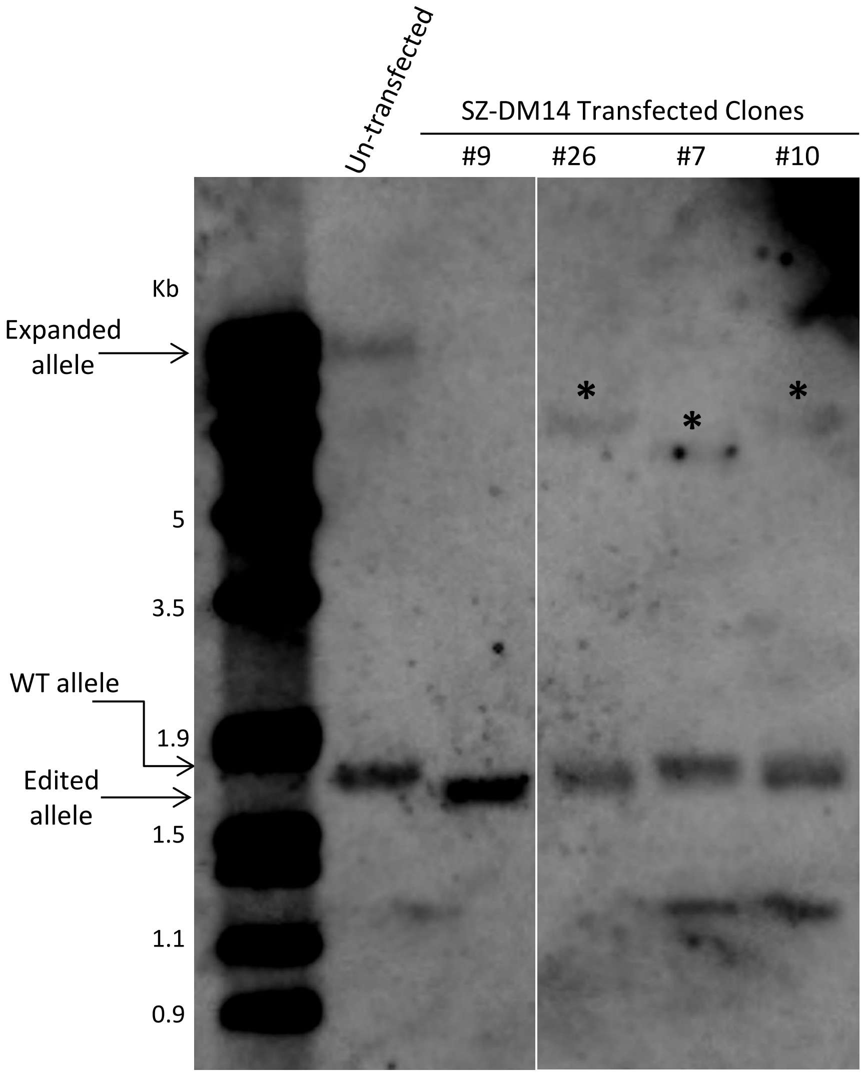

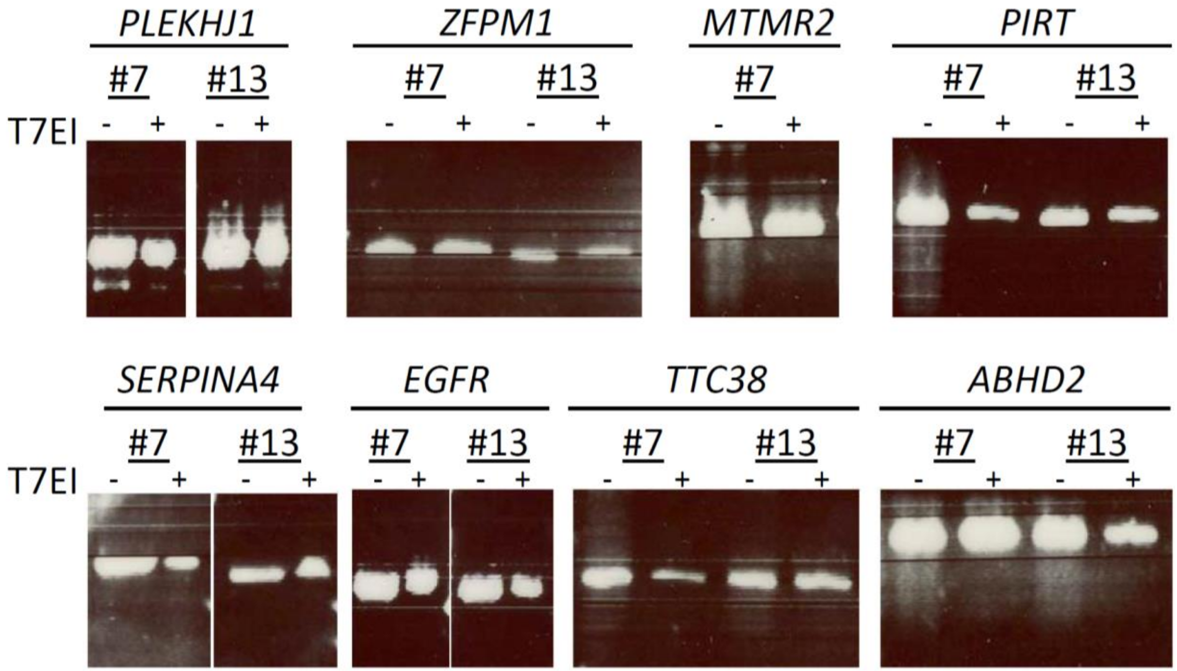

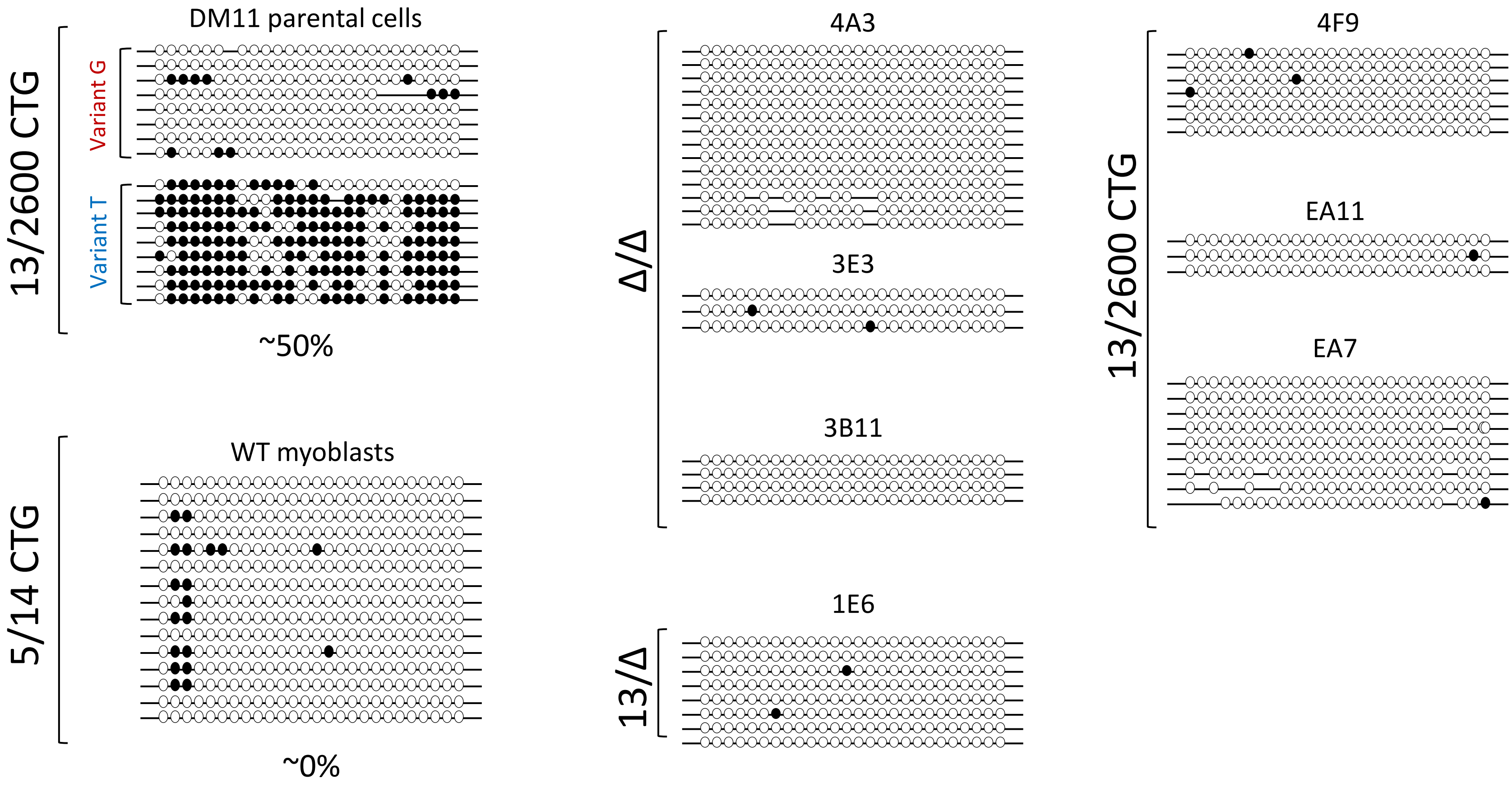
