## Supplementary material for "Deletion of the CTG Expansion in Myotonic Dystrophy Type 1 Reverses *DMPK* Aberrant Methylation in Human Embryonic Stem Cells but not Affected Myoblasts": Tables S1-S3

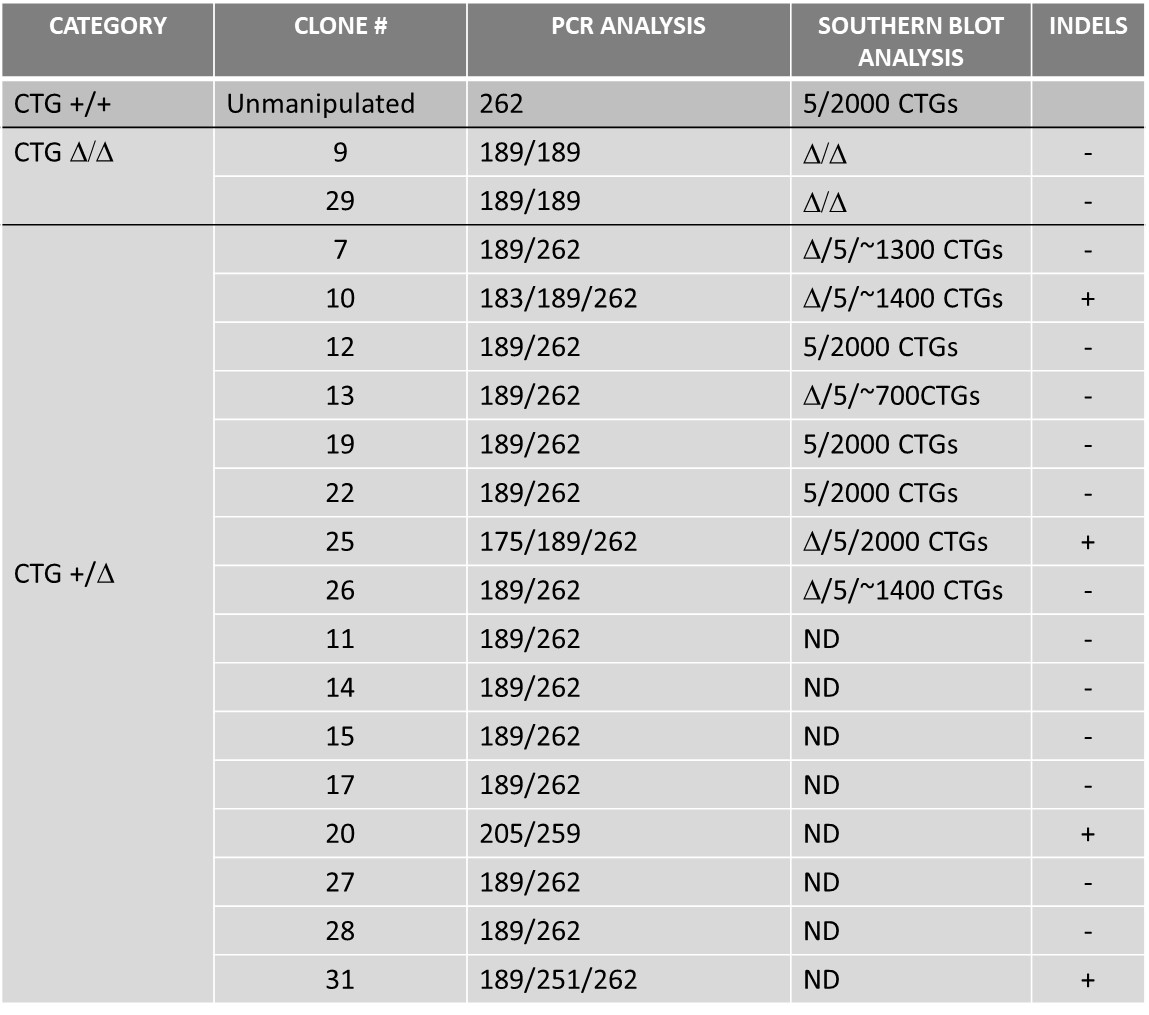

Table S1: Analysis of CRISPR-edited clones of SZ-DM14 hESCs.

27 puro-resistant isolated clones were screened for the expected deletion by Gene Scan analysis of PCR-amplified products from genomic DNA. In 18 clones, successful targeting of the repeat region was identified by the presence of a 189 bp fragment if the normal or expanded alleles were properly edited (loss of 57 bp flanking sequence in addition to 5 or ~2,000 CTG triplets, respectively) in place of or in addition to a 262 bp product from the intact normal allele. PCR products <189 bp were considered as containing indels. Southern blot analysis further validated the deletion of the repeats in a selected number of clones, leaving no doubt regarding the complete removal of the CTG repeat from one/both alleles. Altogether, we achieved 2 clones which were completely *DMPK* repeat deficient with no indels in normal and expanded alleles (Δ/Δ, 7.4%). In the remaining clones (+/Δ), targeting was less efficient and led to a more complex and unpredictable result. The apparent inconsistency in the results between the PCR and the Southern blot assays is most likely due to the difference in sensitivity and quantification ability of these methods. N.D. = not determined.

Table S2: Primers for ChIP analysis by qPCR

| Locus | Forward primer | Reverse primer | Tm | Product size (bp) |
| --- | --- | --- | --- | --- |
| *DMPK* | CTGCCAGTTCACAACCGCTCCGAG | GCAGCATTCCCGGCTACAAGGACCCTT | 60 | 147 |
| *APRT* | GCCTTGACTCGCACTTTTGT | TAGGCGCCATCGATTTTAAG | 60 | 85 |
| *HOXA9* | CTCAGGAGCCTCGTGTCTTT | GTGACCAGGTGGAGGTGTGT | 60 | 82 |

Table S3: Potential off-target sites for 7gRNA and 44gRNA in the human genome.

| Locus Name | On/Off Target Sequence | Primer Sequences |
| --- | --- | --- |
| *DMPK*- 7gRNA target sequence | CAGCAGCATTCCCGGCTACA |  |
| *PLEKHJ1* | CAGAAGCGTTCTCGGCTACA  AGG | F- 5’- ACATAGCGAGACCCCATCAC-3’  R- 5’- GGGACCTGGGACTAGACCAT-3’ |
| *ZFPM1* | CAGCATCCTTCCCAGCTACA  GGG | F- 5’- GTTAATCGCAGCCCTTATCG-3’  R- 5’- TCTGGTTCCTGTCCTTCCAG-3’ |
| *MTMR2* | AAGCAGTCTTCGCGGCTACA  GGG | F- 5’- GCGTAGCCTTCAGAAACCAG -3’  R- 5’- TCCTTATCGCCTTCCTGAGA -3’ |
| *PIRT* | TGGCAACATTCCCGGCTGCA  TGG | F- 5’- GCTAAGGGAGCTAGGGCTGT -3’  R- 5’- GGACTCATGATGCTGGTGTG -3’ |

| Locus Name | On/Off Target Sequence | Primer Sequences |
| --- | --- | --- |
| *DMPK*- 44gRNA target sequence | CAGTTTGCCCATCCACGTCA |  |
| *SERPINA4* | CTGTCTACCCATCCACATCA AGG | F- 5’- CCAAACCAGGACACCAGAGT -3’  R- 5’- CAAGGCCCTGTAGAGGTCAA -3’ |
| *EGFR* | AAGCTTACCCATCCAAGTCA  TGG | F- 5’- TCAGAGGGACAGGAAAGGTG -3’  R- 5’- ATGATTCACAAAGGCGGAAG -3’ |
| *TTC38* | CAGTTTCCCCATCTAAATCA  GGG | F- 5’- TGCTGAGACCTGTTCAGTGC -3’  R- 5’- TGACACATGCCACACTGATG -3’ |
| *ABHD2* | CAGCTTATCCATCCATGTCA  GGG | F- 5’- TTGGTGAACAAGGCAGAGTG -3’  R- 5’- AACTGGACAGAGGGCAAGAA -3’ |

Mismatches between off-target sequences and gRNA sequences are indicated in red, PAM sequence in blue. Primer sequences for each locus are displayed in the right column. Top table for the upstream gRNA (7gRNA) and its predicted off-target and the bottom table for downstream gRNA (44gRNA) and its predicted off-targets.
